# *DICER1* syndrome mutations lead to a gain of 3p-miRNA function, HERVH activity and increased metastatic potential

**DOI:** 10.64898/2025.12.16.694338

**Authors:** Katrina Gordon, Nicolas Bellora, Jeroen Witteveldt, Joanna K. Wojtus, Felix Mueller, Katharina Kases, Pilar G. Marchante, Guillermo Peris, Shelagh Boyle, Sara R. Heras, Atlanta G. Cook, Cei Abreu-Goodger, Sara Macias

## Abstract

The *DICER1* gene is mutated in cancer, including DICER1 syndrome, a rare tumour predisposition syndrome. Cancer-associated hotspot mutations have been reported in both catalytic domains of DICER1 and are predicted to disrupt miRNA biogenesis. To understand how these hotspot mutations contribute to cancer development, we have generated cell lines harbouring single amino acid substitutions within the catalytic RNase IIIa (S1344L) or RNase IIIb (D1709N) domains of the endogenous *DICER1* gene. We show that both mutations result in a widespread loss of 5p miRNAs, and, unexpectedly, an increase in 3p passenger strands loading into AGO2. The shared similarities between both mutants can be attributed to the structural proximity of the S1344 residue to the RNase IIIb catalytic centre. Functionally, we found that changes in the repertoire of miRNAs loaded into AGO2 result in altered gene expression, impacting critical pathways for cancer development, including metastatic potential. Additionally, our results indicate that inactivating the processing activity of DICER1 does not result in genomic instability. Instead, mutations cause specific upregulation of human endogenous retrovirus H (HERVH) through miRNA-independent mechanisms, suggesting that both canonical and non-canonical DICER1 functions are important to understand DICER1 syndrome.

## Introduction

DICER1 is an essential enzyme for the production of microRNAs (miRNAs). In humans, the canonical miRNA biogenesis pathway consists of two processing steps. The first step is nuclear and is carried out by the Microprocessor complex, formed by DROSHA and DGCR8, which generates precursor miRNAs (pre-miRNA). In the second cytoplasmic processing step, DICER1 cleaves the pre-miRNA hairpin to removes the loop structure. This generates a miRNA duplex containing both 5p and 3p miRNAs. Usually, one of the miRNA strands is preferentially loaded into an Argonaute protein (AGO), and named the ‘guide’ strand. The other strand is named the ‘passenger’ and it is usually degraded. This selection seems to be dictated by the thermodynamic stability of the miRNA duplex ends and the 5’ nucleotide identity (reviewed in^1^). *In vitro*, AGO displays preference for 5’ uracil, and preferentially loads the strand with the less stable 5’ end^2^. However, a significant proportion of miRNAs do not follow these rules for strand preference, and selection seems to be context or cell-type dependent^3^. The seed sequence, in positions 2 to 8 relative to the 5’end of the miRNA, is critical for directing the RNA-induced silencing complex (RISC), containing AGO loaded with miRNAs, by complementary sites in the 3’ untranslated region (UTR).

While insects, nematodes and plants possess duplicated *DICER* genes, with specialised paralogs for miRNA biogenesis or the production of small interfering RNAs (siRNAs), mammals only express a single *Dicer* gene that needs to be able to process both miRNA and siRNA precursors (reviewed in^4^). Although it is still unclear if mammals possess a functional endogenous siRNA (endo-siRNA) pathway, miRNA-independent functions of mammalian Dicer have been reported. These include the maintenance of genomic stability and proper chromosome segregation, as well as preventing the accumulation of transcripts derived from the major satellite repeats from pericentromeric regions^5–11^. In addition, Dicer has also been implicated in regulating other types of repeats, including retrotransposons (reviewed in^12,13^). In support of this, Dicer was found to repress the expression of LTR and LINE-1 retrotransposons during early mouse development and mouse embryonic stem cells (mESCs)^14–17^. Although the mechanisms driving transposon silencing remain unclear, LINE-1 transcripts are predicted to generate siRNAs with silencing activity. The LINE-1 5’UTR contains a bidirectional promoter, presumably resulting in the formation of dsRNA that can act as a substrate for Dicer cleavage^18^. Other endo-siRNAs have been detected in mouse oocytes and mESCs. These are predicted to regulate the main retrotransposon families, LINE, SINE and LTR^19–21^.

DICER1 mutations have been identified in tumours associated with DICER1 syndrome. This syndrome increases the predisposition to both benign and malignant tumours. Patients typically carry an inherited loss-of-function mutation in one *DICER1* allele and acquire a second somatic mutation later in life. While the loss-of-function germline mutations can occur in any region of the protein, the somatic mutations usually accumulate within the catalytically active, metal-ion binding residues^22^. These include amino acid substitutions in both RNase IIIa and IIIb endonucleolytic domains. The RNase IIIa domain cleaves the 3p-arm within the pre-miRNA hairpin, while the RNase IIIb cleaves the 5p-miRNA (reviewed in^23^). Despite being in different RNase III domains, most cancer-associated mutations are predicted to only affect 5p miRNA production. Intriguingly, there does not seem to be a selective advantage for mutations affecting 3p miRNA production in cancer^24^.

Considering that DICER1 is an essential protein for cell survival, the observation that inactivating its catalytical activity can contribute to cancer is counterintuitive^25,26^. We have generated human cells recapitulating cancer-associated hotspot mutations using CRISPR/Cas9 editing. By comparing different mutations in the RNase IIIa and IIIb domains, we have confirmed that mutations in either domain results in inactivation of 5p miRNA production. However, unexpectedly, there is a concomitant increased loading of many 3p miRNAs into AGO2, leading to numerous cases of miRNA arm switching. This results in defective silencing of 5p-miRNA targets, and increased repression of 3p-target mRNAs. Our results show that the hotspot mutation S1344L in the RNase IIIa also affects 5p-miRNA cleavage, but to a lesser extent than RNase IIIb D1709N. Analysis of DICER1 structures show that S1344 is spatially close to the RNase IIIb active site, supporting its role in 5p-miRNA processing. Mechanistically, we show that DICER1-associated mutations do not cause genome instability or chromosome abnormalities, as suggested for *Dicer* knockout cells. Instead, these mutations are associated with upregulation of endogenous retroviruses (ERVs) associated with a ‘pluripotency’ profile. Interestingly, despite having a phenotype of defective growth, DICER1-mutant cell lines showed increased metastatic potential, providing an explanation of how these mutations can predispose to cancer progression.

## Results

### Generation of DICER1-syndrome mutation cell lines

To study the role of DICER1-syndrome associated hotspot mutations, we generated human cell lines containing single-amino acid substitutions in *DICER1* using CRISPR/Cas9. We used PA-1 cells, a diploid human embryonic cell line derived from an ovarian teratocarcinoma. We introduced the S1344L substitution in the RNase IIIa domain, and the D1709N substitution at the RNase IIIb domain, which are typical hotspot mutations found in DICER1-syndrome patients (**Figure 1A**). Cell lines where both alleles were modified with the same mutations (S1344L or D1709N) were named homozygote mutants (HOM). We also generated compound heterozygote (HET) mutants. HET cell lines contained only one allele mutated either as S1344L or D1709N, while the second allele contained an indel mutation generating a premature-termination codon (PTC), driving the resulting mRNA to be degraded by non-sense mediated decay (NMD) (**Figure 1B**). The HET cell lines are more representative of the genetic composition observed in DICER1 syndrome patients, where one allele contains a loss-of-function mutation that results in loss of protein production, while the second allele acquires a hotspot mutation in RNase IIIa or IIIb sites. No knock-out cell lines were obtained for the *DICER1* gene, suggesting that in human embryonic teratocarcinoma cells, DICER1 is essential for viability as observed in human embryonic stem cells ^26^.

**Figure 1.**
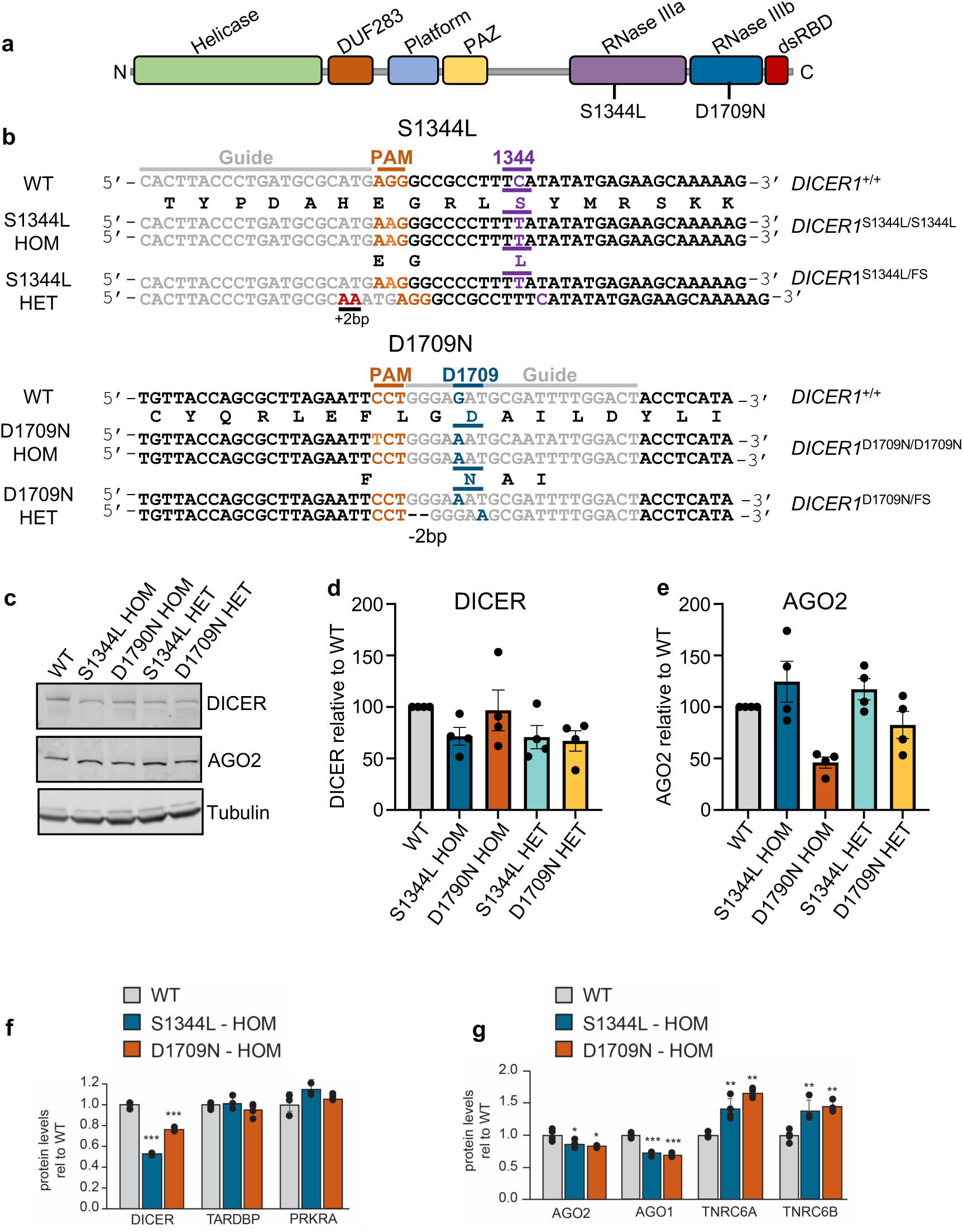
Generation of DICER1-mutant PA-1 cell lines harbouring point mutations in either RNase IIIa or RNase IIIb domain. (**a**) Schematic of DICER1 protein showing functional domains as indicates (DUF: domain of unknown function). Targeted point mutations at amino acid residue S1344L and D1709N in the RNase IIIa and RNase IIIb domains are shown. (**b**) CRISPR/Cas9 knock-in strategy showing the sequence of the targeted region in PA-1 wild type (WT) cells and the corresponding sequence changes in each of the mutant clones as indicated with amino acid change S1344L (purple) and D1709N (blue), Guides (grey) and PAM (orange). Both alleles are shown for each of the mutants, homozygous (S1344L HOM and D1709N HOM) and frameshift (FS) allele are indicated in the compound heterozygous lines (S1344L HET and D1709N HET). Genotypes are as follows: S1344L HOM (*DICER1*^S1344L/S1344L^), S1344L HET (*DICER1*^S1344L/FS^), D1709N HOM (*DICER1*^D1709N/D1709N^) and D1709N-HET (*DICER1*^D1709NL/FS^). (**c**) Western blot analysis of DICER1 and AGO2 protein levels in WT, S1344L and D1709N mutant cell lines with tubulin as a loading control (**d** and **e**) Quantification of DICER1 (**d**) and AGO2 (**e**) protein levels relative to WT. Error bars indicate ± S.D. from three independent experiments. (**f** and **g**) Proteomic data analysis of DICER1 binding partners TARDBP and PRKRA (**f**) and components of the miRISC complex, AGO1/2 and TNRC6A/B (**g**) in WT and HOM mutants from total cell extracts. Data are presented as the average ± S.D. from four independent experiments. Error bars indicate ± SD and significance is indicated as *** p ≤ 0.001, ** p ≤ 0.01, * p ≤ 0.05, ns (not significant) p ≥ 0.05 by one-way ANOVA followed by Tukey’s multiple comparison test.

We first assessed DICER1 and AGO2 protein levels by western blotting (**Figure 1C**). Interestingly, no significant differences were observed between HET and HOM mutants, despite HETs only producing DICER1 protein from a single allele (**Figure 1D and 1E**). In agreement, *DICER1*^+/-^ PA-1 cells, where only a single allele is inactivated, do not display significant changes in miRNA expression, arguing against DICER1 haploinsufficiency (**Suppl Fig 1).** Proteomics data confirmed that DICER1 mutations did not result in significant changes in the protein levels of its co-factors, TARDBP or PACT (PRKRA) (**Figure 1F**), while AGO2 and AGO1 displayed significantly decreased levels (**Figure 1G**). Other components of the RNA-inducing silencing complex, such as the GW182 proteins (TNRC6A and B) were not decreased by DICER1 mutations (**Figure 1G and Suppl. Excel File 1**).

### Increased cell migration in DICER1-syndrome cells

Considering that *DICER1* is an essential gene in both mouse and in human cells, it is unclear how its mutation relates to tumorigenesis. To explore the mechanisms by which DICER1 mutation predisposes to cancer, we decided to explore different phenotypes of the generated cell lines. In agreement with its reported role in cell survival and cell cycle progression, we confirmed that all *DICER1* mutant cells showed significantly lower replication rates compare to wild-type (WT) cells (**Figure 2A and Suppl Fig 2A**). These mutations also resulted in a decreased capacity for colony formation (**Figure 2B-2C and Suppl Fig 2B**). On the other hand, mutant DICER1 cell lines display increased metabolic activity (**Figure 2D**), and an increased ability to migrate in a wound healing assay (**Figure 2E and 2F**). To investigate which gene expression changes supported these phenotypes, we performed differential gene expression analyses of total RNA high-throughput sequencing in both S1344L and D1709N cell lines compared to WT. Interestingly, D1709N cells showed more significantly expressed genes than S1344L, but there was a clear overlap between the two mutants (**Suppl Excel File 2**). More than 80% of the genes that were differentially expressed in the S1334L mutant cell line were also differentially expressed in D1709N cells. Considering that both mutations result in similar cellular phenotypes, we decided to focus our analyses on the commonly dysregulated genes. Genes upregulated upon DICER1 mutations (N=1,306) were associated with ‘Epithelial to mesenchymal transition (EMT)’ and ‘KRAS signalling’ (**Figure 2G**). These pathways agree with the increased migration potential of *DICER1* mutant cells in the wound healing assay, as EMT is associated with tissue repair, but also increased motility, invasiveness, metastasis and enhanced resistance to apoptosis ^27,28^. The contractile properties of cells during migration may explain enrichment of the ‘myogenesis’ pathway with the upregulated genes. KRAS signalling is also associated with increased invasion and metastasis, and constitutive activation of KRAS is observed in tumours^29^. On the other hand, commonly downregulated genes (n=1,174) were associated with ‘oxidative phosphorylation’ and ‘MTORC1 signalling’ (**Figure 2H**). These pathways may be relevant to the defects we observed in cell growth and alterations in the metabolic state that are also associated with EMT and the Warburg effect in cancerous cells^28,30^. All these findings suggest that despite the critical role for DICER1 in cell survival, DICER1-syndrome associated mutations cause increased cell migration, thus explaining the role of this gene in cancer progression, and possibly metastatic potential.

**Figure 2.**
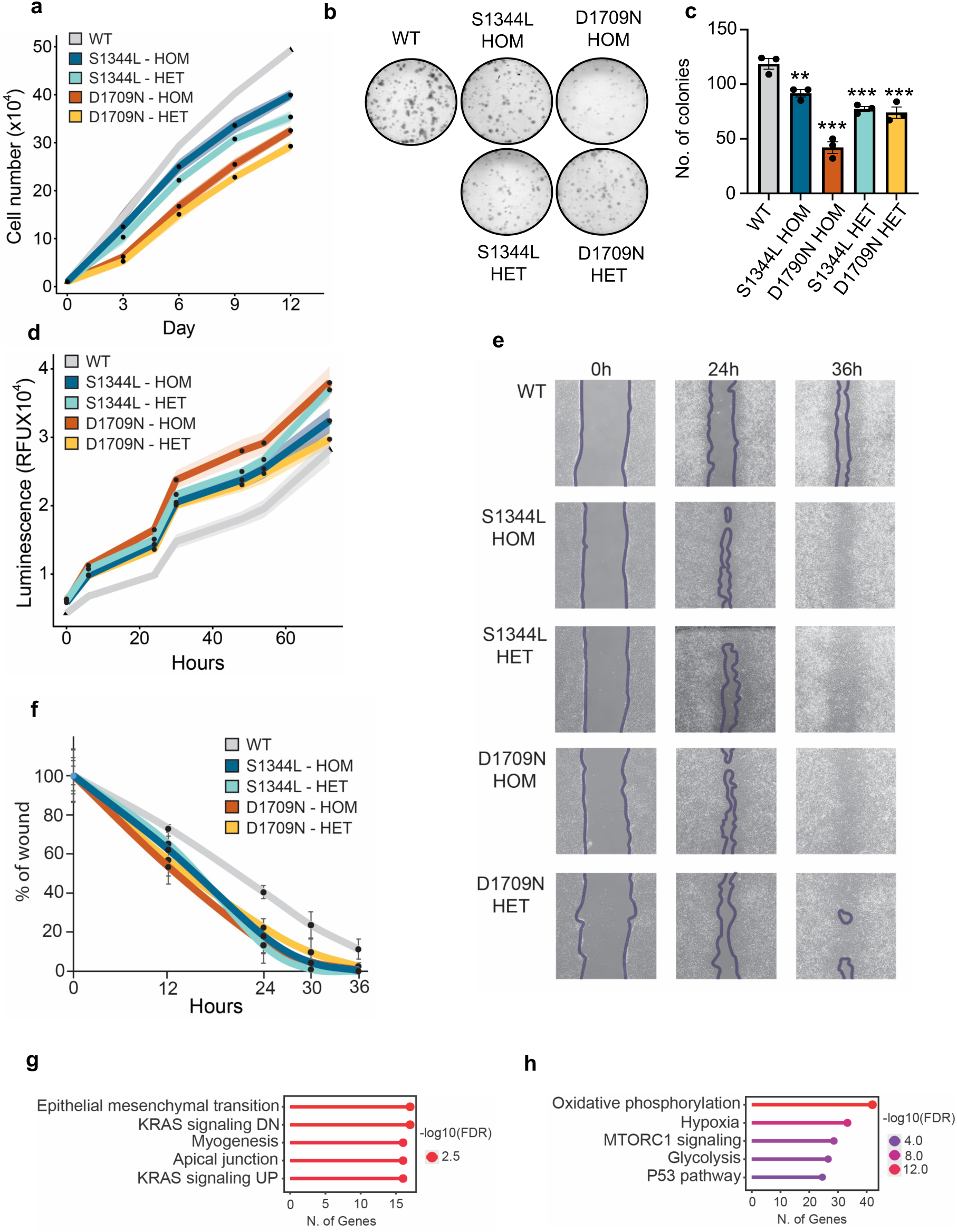
Differences in cellular phenotypes in the DICER1-mutant cell lines. (**a**) Proliferation rates of the four DICER1 mutant cell lines were compared to the parental WT PA-1 cells. Shaded lines indicate ± SD of three independent replicates. (**b**) Representative images of crystal violet-stained colonies indicating the colony formation capacity of each of the cell lines. (**c**) Quantification of the number of colonies. Error bars indicate ± SD of three independent replicates. Error bars indicate ± SD and significance is indicated as *** p ≤ 0.001, ** p ≤ 0.01, * p ≤ 0.05, ns (not significant) p ≥ 0.05 by one-way ANOVA followed by Tukey’s multiple comparison test. **(d)** Metabolic activity across the different cell lines was assessed by quantifying cellular ATP via a luciferase reaction and measuring the resulting luminescence over time. Shaded lines indicate ± SD of four independent replicates. (**e)** A wound healing assay was used to compare the migratory potential of the different cell lines. Representative phase contrast images are show at time 0 hrs, 24 hrs and 36 hrs after initial wound. (**f**) Quantitative analysis of wound closure expressed as a percentage of the original wound. Data are presented as the average ± SD from three independent experiments (n=3). All mutants (except S1344L-HET at t=12h) have significantly higher rates of wound closure at all time points compared to WT (p<0.007). Data was analysed by one-way ANOVA, followed by a Tukey HSD/Tukey Kramer post-hoc test. (**g-h**) Hallmark gene sets for the commonly differentially expressed up (**g**) and down (**h**) regulated genes in DICER1 mutants compared to WT PA-1 cells (only the top5 gene sets are shown).

### S1344L and D1709N result both in 5p-miRNA loss and 3p-miRNA gain

To understand the mechanism by which mutations in DICER1 syndrome caused these altered cellular phenotypes, we initially assessed their effects on miRNA production. We measured miRNA levels in cells using two alternative approaches, small RNA-sequencing and high-throughput RNA sequencing of AGO2 immunoprecipitate (IP). By comparing these two datasets, we can firstly quantify which miRNAs are differentially expressed, and secondly, which miRNAs are differentially loaded into AGO2, driving changes in post-transcriptional gene silencing. Our results showed that despite being in two distinct RNase III domains, both S1344L and D1709N mutations resulted in decreased abundance of 5p-miRNAs in both the small RNA and AGO2 IP datasets, relative to WT cells (**Figure 3A and 3B**, left column **and Suppl Excel File 3**). This suggests that both mutations have a similar effect on DICER1 activity. We note that despite being in the RNase IIIa domain, the S1344 residue of DICER1 is structurally close to the catalytic site responsible for 5p-miRNA cleavage (**Figure 3C and 3D**)^24^. On analysis of an experimentally determined structure of human DICER1 in its dicing state shows that S1344 forms a hydrogen bond with the 2’OH of the 23^rd^ base from the 5’ end, orienting the backbone phosphate group linking the 22^nd^ and 23^rd^ nucleotides close to the RNase IIIb catalytic divalent cation^31^. When modelling these mutations *in silico*, substituting the serine with leucine in position 1344 generates a steric clash with the pre-mRNA backbone near the active site of the RNase IIIb domain and so may prevent correct positioning of the 23rd phosphate group in the active site (**Figure 3D**). On the other hand, the divalent cation in the active site is coordinated by four acidic side chain (E1705, D1709, D1810 and E1813) that are all known hotspot sites for DICER1 mutations^24^. The D1709N mutation neutralizes the negative charge at this residue position, which likely interferes with the catalytic metal ion binding to the enzyme. The difference in structural consequences of mutating the D1709 vs S1344 position explains why D1709N displays a more pronounced 5p miRNA loss than S1344L.

**Figure 3.**
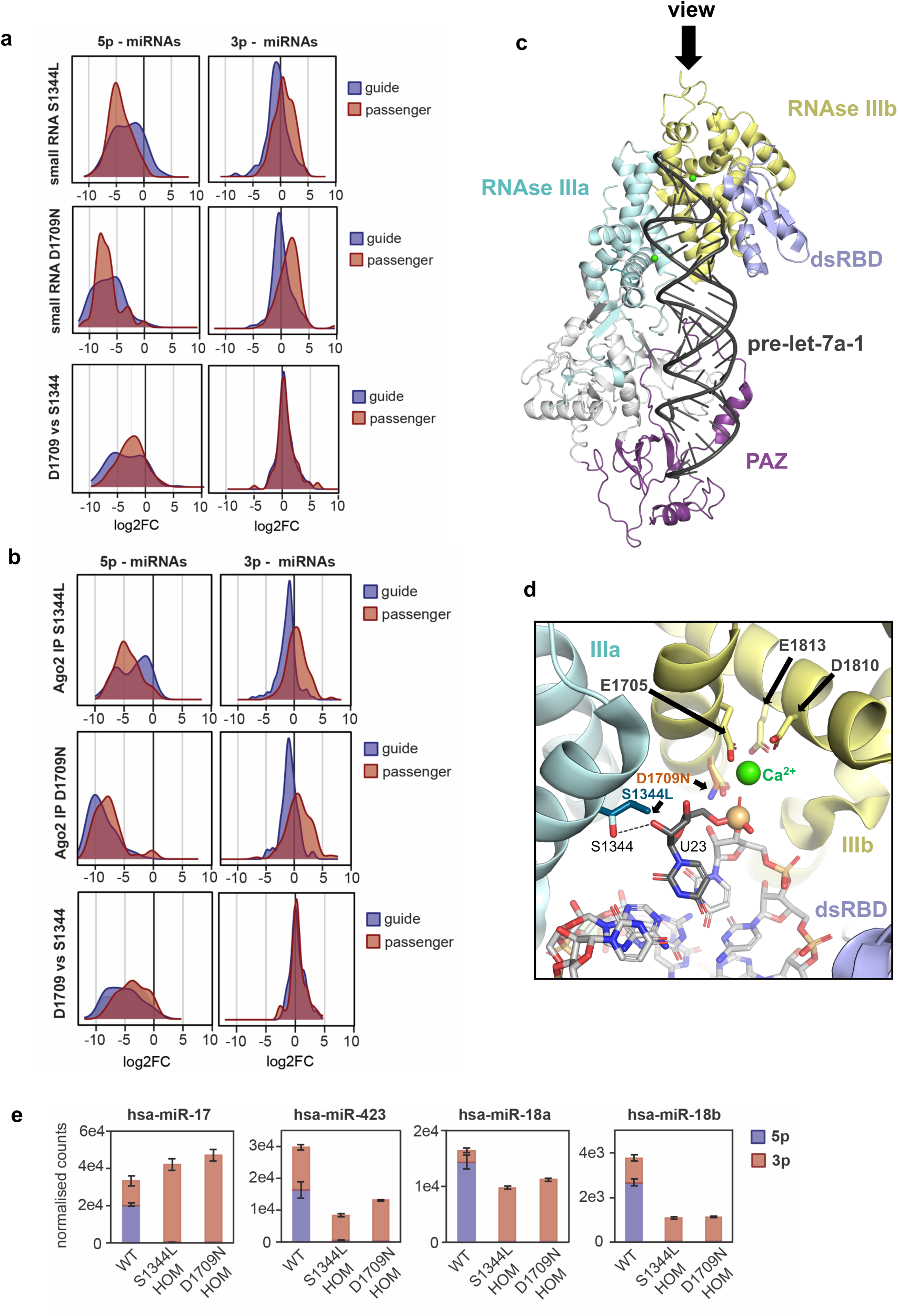
Accumulation of 3p-miRNAs in DICER1 mutated cell lines. **(a)** Density plots showing the change in distribution of 5p and 3p miRNAs in the mutants relative to WT and to each other using the small RNA sequencing datasets. Guide strand (red) and passenger strand (blue). For definition of guide please see Suppl Figure 3a. **(b)** Density plots showing the change in distribution of 5p and 3p miRNAs in the mutants relative to WT and to each other using the AGO2-IP dataset. Guide strand (red) and passenger strand (blue). **(c)** Overview of pre-let-7a-1 bound to human DICER1 in the dicing state (PDBid 7XW2). PAZ (violet), RNase IIIa (cyan), RNase IIIb (yellow) and dsRNA binding domain (dsRBD, light blue) are highlighted. (**d**) A viewpoint taken from the top of the pre-miR helix, showing the active site of RNase IIIb. A divalent cation (Ca^2+^, green) in the active site is coordinated by four acidic side chain (E1705, D1709, D1810 and D1813). Ser1344 interacts (black dotted line) with the ribose 2’OH of U23. The phosphate group of U23, which is the site of hydrolysis, is highlighted with an orange sphere at the phosphate atom. Superimposed is the leucine sidechain (teal) of a S1344L mutant and the asparagine sidechain (orange) of D1709N. (**e**) Examples of miRNA strand switch in DICER1 mutant cells. Data are represented as the average of normalized counts +/- SD. 5p miRNA in blue, and 3p miRNA in red.

Unexpectedly, mutations in DICER1 endonucleolytic activity also resulted in a gain of some 3p-miRNAs, in both the small RNA and AGOP2 IP dataset. First, we found that highly abundant 3p miRNAs in WT cells, such as miR-302-3p, were not upregulated in comparison with other less abundant 3p miRNAs (**Suppl Excel File 3**). We then investigated whether these differences were dictated by the initial abundance of the 3p miRNA, being an abundant guide, or alternatively a less abundant passenger. To test this possibility, we first classified all 5p and 3p miRNAs into guides or passengers depending on their abundance in WT cells (**Suppl Fig 3A**). MiRNAs with similar abundance for the 5p and 3p strands were excluded from this analysis. We next plotted the differences in expression of 3p guide and passenger miRNAs, and whilst 3p guides did not change expression or were slightly reduced in DICER1 mutants, 3p passengers showed increased abundance in both the small RNA and the AGO2 IP datasets (**Figure 3A and 3B**, right column). Both the S1344L and D1709N mutants behaved similarly, suggesting that this defect is not the result of increased processing of the 3p miRNA, but increased loading efficiency into AGO2, in cases where the 5p miRNA guide cannot be loaded. Two recent independent studies have shown increased AGO2 loading of 3p-passenger miRNAs in the context of *DICER1* mutations^32,33^. In agreement with our findings in cell models, TCGA tumor datasets with homozygous *DICER1* mutations also showed decreased 5p-miRNA levels and increased 3p-miRNA abundance (**Suppl. Fig 3B**). We conclude that DICER1 mutations cause a general loss of 5p miRNAs in addition to a gain-of-function of 3p miRNAs that are not usually expressed in healthy or WT cells. In some extreme cases, this results in a complete miRNA strand switch, suggesting a shift in the mRNAs being silenced (**Figure 3E**).

The unexpected accumulation of 3p-miRNAs led us to investigate if this could be contributing to alterations in post-transcriptional gene silencing. To this end, we analysed the cumulative distribution function (CDF) of genes predicted to be targeted by the dysregulated 3p and 5p-miRNAs. This analysis revealed that targets of the increased 3p miRNAs were destabilised in D1709N cells, both at the transcriptomic and proteomic level (**Figure 4A and 4B**). On the other hand, targets of the 5p-miRNAs increased their steady state levels, as 5p-miRNAs were globally decreased in D1709N mutant cell lines, again at the transcriptomic and proteomic level. Using the same approach, we measured the effect of the S1344L mutation. In this case, no significant changes were observed, except for the 5p-miRNA targets at the proteome level (**Figure 4C and 4D**), confirming again that the molecular defects observed by S1344L mutation are less pronounced than D1709N mutants.

**Figure 4.**
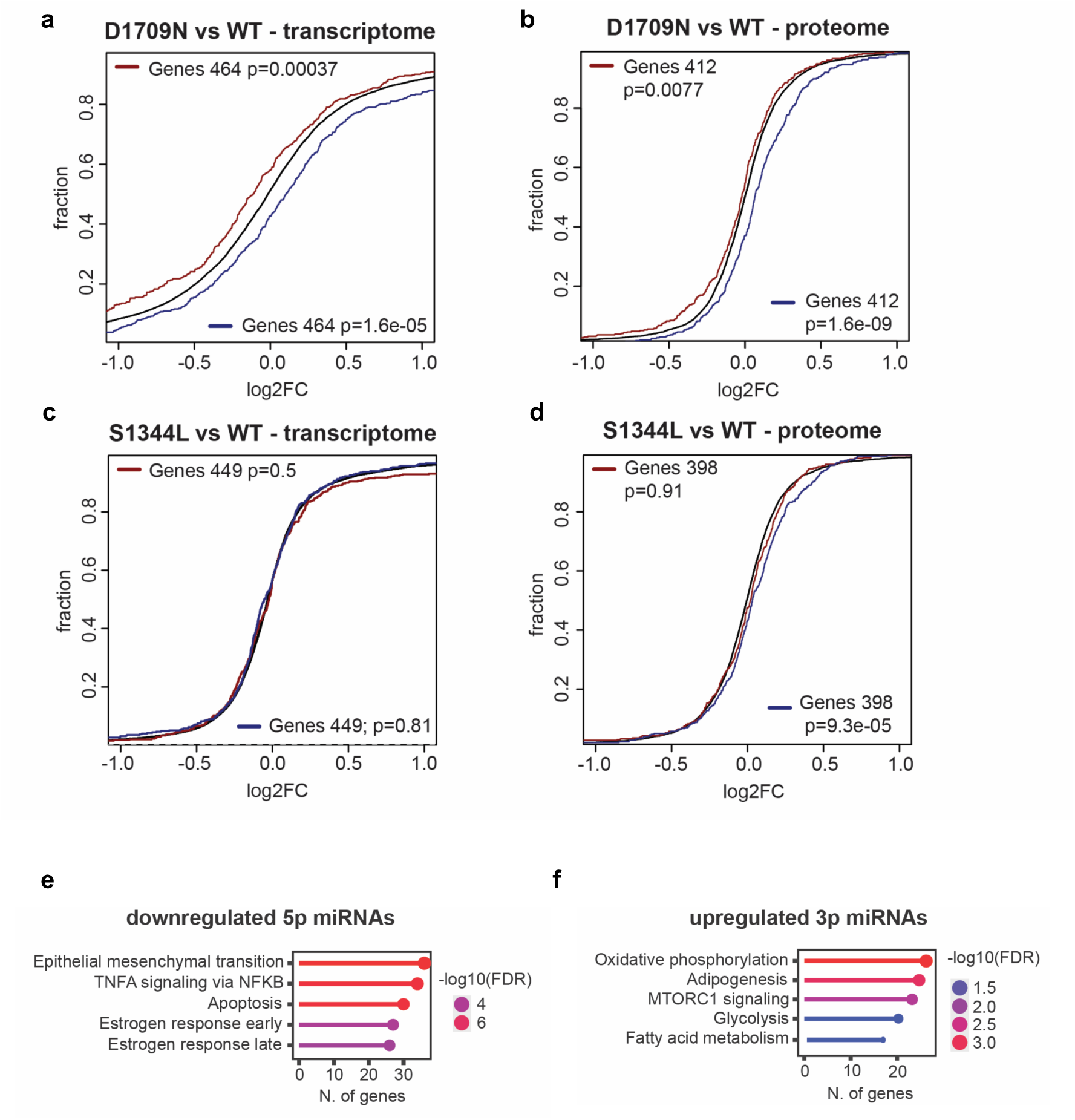
Impact of 3p and 5p-dysregulation in DICER1 mutated cell lines. Cumulative distribution function (CDF) vs log2 fold-changes of predicted gene targets for upregulated (red) and downregulated (blue) miRNAs in the D1709N mutant compared to WT, measured by total RNA-seq (**a**) or total cell proteomics (**b**). Cumulative distribution function (CDF) vs log2 fold-changes of predicted gene targets of upregulated (red) and downregulated (blue) miRNAs in the S1344L mutant compared to WT, measured by total RNA-seq (**c**) and total cell proteomics (**d**). p-val indicates if log2 fold-changes (log2FC) of target genes are significantly down (red) or upregulated (blue). (**e**) Hallmark gene sets of the predicted gene targets for the top 10 downregulated miRNAs (5p) which were significantly increased in the transcriptomic data (log fold 2 > 0.5, p-adj. < 0.05). (**f**) Hallmark gene sets of the predicted gene targets for the top 10 upregulated miRNAs (3p) which were significantly decreased in the transcriptomic data (log fold 2 > 0.5, p-adj. < 0.05).

Finally, we explored the pathways associated with the predicted targets for the differentially expressed 5p and 3p miRNAs. We only focused on those targets with significant changes in expression in the RNA-seq dataset. Interestingly, the predicted targets of the downregulated 5p miRNAs were involved in epithelial to mesenchymal transition (EMT) (**Figure 4E**). On the other hand, the predicted targets of the upregulated 3p miRNAs were associated with metabolic pathways (**Figure 4F**). It seems that both 5p and 3p miRNA dysregulation can contribute to defective post-transcriptional gene silencing in DICER1 mutant cells, and that these changes may contribute to the altered cellular behaviours observed in wound healing and metabolism.

### Dysregulated ERV profile in DICER1-syndrome cell models

In addition to miRNA production, mammalian DICER1 is known to be involved in maintenance of genome stability as well as chromosome segregation during cell division^5,6,9–11^. Most of these studies have been performed in cells where DICER1 protein is absent, but the role of the endonucleolytic activity of DICER1 on these functions has not yet been explored. To study genome stability, we performed metaphase spreads of all the DICER1-mutated PA-1 cell lines. PA-1 cells are a female diploid human embryonic teratocarcinoma cell line (46, XX), and all mutants displayed the same chromosome count, indicating that *DICER1* mutations are not associated with increased genome instability or improper chromosome segregation (**Figure 5A**).

**Figure 5.**
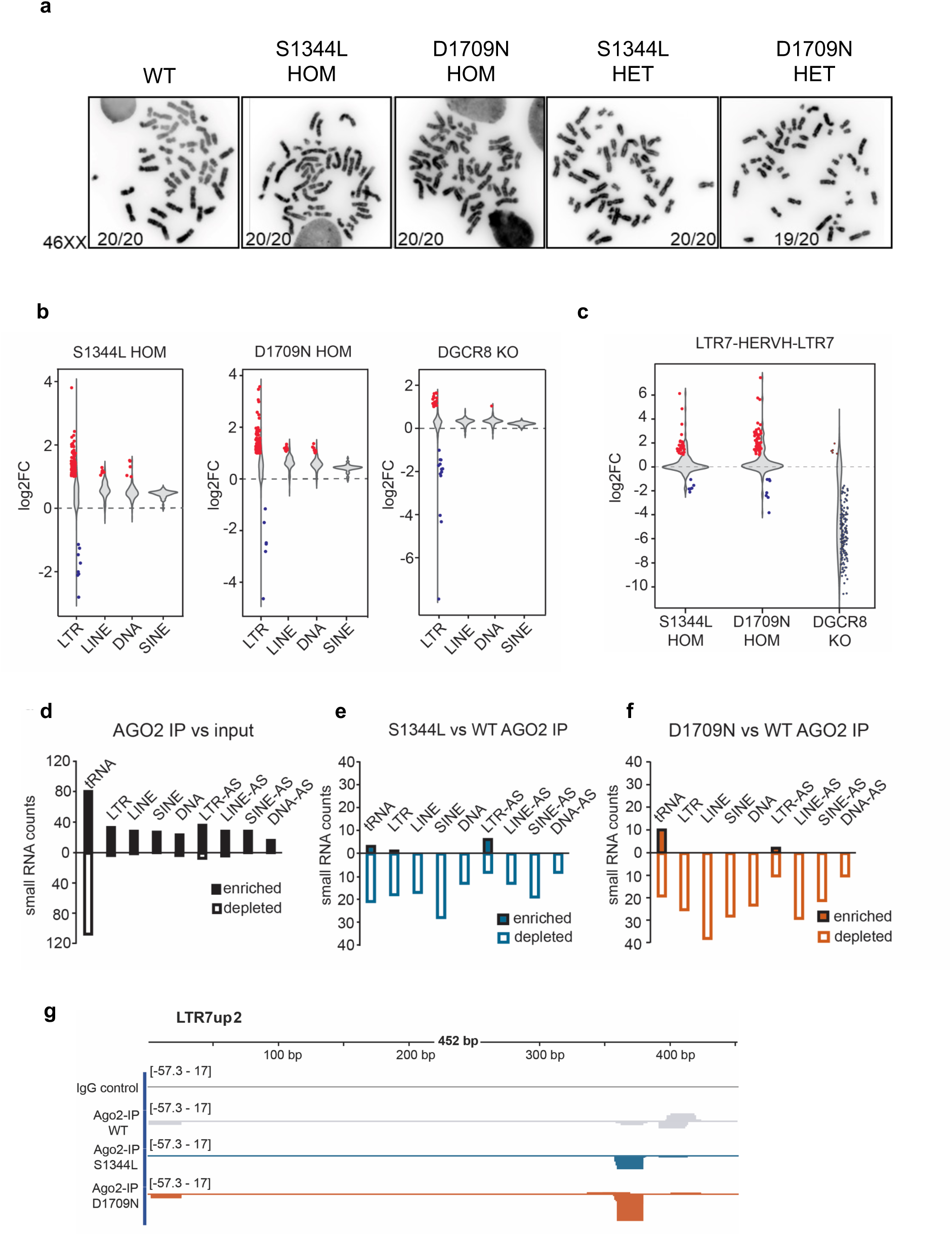
DICER1 mutations and genome integrity. **(a)** Representative images of metaphase spreads (n=∼20) from parental PA-1 (WT) and the four DICER1 mutant cell lines. Counts represent number of spreads with 46XX chromosomes **(b)** Violin plots for differential expression analyses of LTR, LINE, SINE retrotransposons and DNA transposons, in S1344L HOM (left), D1709N (middle), and DGCR8 KO (right) PA-1 cells versus WT. Significantly upregulated families (red), and downregulated (blue) are shown as individual dots. **(c)** Violin plots for differential expression analysis of defragmented LTR7-HERVH-LTR7 elements. Each dot represents a unique locus, of significantly upregulated (red) and downregulated (blue) loci. (**d**) Enrichment of small RNA counts mapping to sense and antisense (AS) LTR, LINE, SINE and DNA elements in the AGO2 immunoprecipitations in PA-1 WT cells versus the small RNA input (**e**) Differential enrichment analysis for small RNAs mapping to repeats in the S1344L AGO2 IPs vs WT cells. (**f**) the same as (e) for the comparison of D1709N AGO2 IP vs WT (**g**) Mapping of AGO2 IP-associated small RNAs to the LTR7up2 consensus sequence. Samples included are the IgG control, AGO2 IP from WT cells (grey), from S1344L cells (blue), and D1709N cells (orange), in both in the sense (+) and antisense (-) strand.

Mammalian DICER1 has also been proposed to be involved in the generation of endo-siRNAs or small RNAs with silencing properties against repetitive sequences^12^. To test if this function could be relevant in DICER1-syndrome, we first assessed transposable element (TE) expression in DICER1 mutant cells compared to WT using the total RNA high-throughput sequencing datasets. For this purpose, we used an in-house annotation of TEs, in which fragmented elements were merged into full-length copies (see Methods). These analyses revealed that both S1344L and D1709N mutations resulted in a general derepression of all types of TEs, i.e. both retrotransposons (including LTR, LINEs and SINEs) and DNA transposons (**Figure 5B**). LTRs, including endogenous retroviruses, such as the ERV1 subfamilies, displayed the most dramatic changes (**Suppl Fig 5A-B**). To determine whether this was an indirect effect driven by alterations in miRNA expression, we measured TE expression in *DGCR8* knockout PA-1 cells, as these lack most canonical miRNAs^34^. This comparison revealed that *DGCR8* loss did not result in such a profound general upregulation of TEs as in the *DICER1* mutant cells (**Figure 5B and Suppl Fig 5C**). More detailed analysis revealed that for some TE types, *DGCR8* and *DICER1* mutant cells resulted in opposite behaviours. For instance, the human endogenous retrovirus H (HERVH) was specifically upregulated in *DICER1* mutant cells but not *DGCR8*, strongly suggesting a miRNA-independent regulation for this TE (**Figure 5C and Suppl Fig 5C**). Although TEs are generally silenced in somatic adult cells, HERVH is typically expressed in human early development, and it can be subdivided into several classes depending on the LTR7 sequence^35^. Intriguingly, the most upregulated subfamily in *DICER1* mutant cells were the LTR7up1/2 (**Suppl Fig 5D**), which are highly expressed in pluripotent cells and required to maintain pluripotency^36^.

We next explored the mechanisms by which DICER1 could suppress TE expression. Our results suggested that this regulation was miRNA-independent, although we cannot totally rule out this possibility as DICER1 mutant cells also display miRNA expression changes that may influence TE expression. Alternatively, TE dysregulation could be explained by the miRNA-independent roles of DICER1, including the generation of small RNAs or siRNAs against repeats thus silencing their expression. To test this possibility, we first assessed if AGO2 was associating with small RNAs derived from TEs. Here, we compared small RNAs mapping to repeats in AGO2 IP versus the input (small RNA dataset). We confirmed that small RNAs mapping to both sense and antisense strands of all types of transposons were detected, including LTR, LINE, SINE and DNA, while tRNAs were equally enriched and depleted, indicating a non-specific association with AGO2 (**Figure 5D**). Next, we compared if the association of these small RNAs was altered in *DICER1* mutant cells by comparing the AGO2 IP datasets from WT vs S1344L and D1709N cells. This analysis revealed a reduction in loading of both sense and antisense small RNAs against all types of TEs in both *DICER1* mutant cell lines (**Figure 5E and 5F**). More detailed analysis of specific LTR7 subfamilies, such as the LTR7up2, revealed that *DICER1* mutant cell lines altered the proportion and composition of sense and antisense small RNAs associated with AGO2 that could potentially regulate the expression of LTR7 transcripts (**Figure 5G**). All these together suggest that mutations of the endonucleolytic function of DICER1 alter loading of sense and antisense small RNAs that are complementary to transcripts derived from TEs, regulating their post-transcriptional gene silencing.

Altogether, we suggest that both the functions of mammalian DICER1 in miRNA biogenesis, as well as its function in controlling TE expression are critical to understand the molecular mechanisms by which DICER1 mutations can cause DICER1-syndrome.

## Discussion

Mutations in the *DICER1* gene are associated with the DICER1 cancer predisposition syndrome. However, DICER1 is not the only small RNA biogenesis factor that has been found to be mutated in cancer. DICER1 along with a number of key components of the small RNA biogenesis pathways, such as DROSHA and DGCR8, are found to be mutated in Wilms tumours, the most prevalent paediatric kidney malignancy^37–40^. Similar to our DICER1-mutated cell models, DROSHA mutations are also associated with increased mesenchymal phenotypes^41^. However, in tumours, DROSHA is typically only mutated in a single allele that behaves as a dominant negative. Often, these hotspot mutations affect the RNase IIIb domain, disrupting the 5’end cleavage of the pri-miRNA^38,40^. In contrast, mutations in DGCR8 usually affect both alleles and interfere with its dsRNA binding activity, leading to impaired pri-miRNA processing^37,39,41^. Given that DGCR8 and DROSHA function upstream of DICER1, mutations in these factors result in unproductive processing, leading to depletion of both 5p and 3p miRNAs. This phenomenon has been verified in cell models and tumours carrying DROSHA mutations^38,39,42^.

The specific loss of 5p miRNAs upon DICER1 mutations had been reported in earlier studies^42,43^. While initial research suggested that 3p miRNAs were not affected, more recent studies observed an increase in 3p miRNA abundance in tumours^24,44,45^. Only recently has this increase in 3p miRNA been confirmed in cell models recapitulating DICER1 syndrome mutations^32,33^. Jee *et al* and Malagobadan *et al* confirmed the increased abundance of 3p passenger miRNA loading into AGO2 using a mutant of the RNase IIIa in human stem cells and a mutant of the RNase IIIb in mouse cells, respectively^32,33^. However, a direct comparison between the consequences of human RNase IIIa and IIIb domain hotspot mutations within the same genetic background was not performed in these studies. Our results indicate that whilst both mutations result in a similar increase of 3p miRNAs, the D1709N mutation causes a more pronounced loss of 5p miRNAs compared to S1344L. This is consistent with the structure of human DICER1 where both residues contribute to placing the dsRNA substrate in the correct position in the active site of the enzyme^31^. While S1344 is structurally part of the active site for 5p miRNA, unlike D1709, it does not participate in metal-ion binding required for catalysis, suggesting that some (low) level of 5p processing might still be possible in these mutants.

Supporting the relevance of 3p-miRNA dysregulation in DICER1 syndrome tumours, our findings demonstrate that the expression of predicted targets for these miRNAs is significantly altered. This suggests that, unlike mutations in DROSHA and DGCR8, the molecular phenotypes associated with DICER1 mutations originate from both the loss and gain of specific miRNAs. It will be interesting to explore whether tissues commonly affected by DICER1 mutations, such as the pleura in the lungs, ovaries, kidneys or thyroid, exhibit a bias towards 5p or 3p guides^46^. For instance, are tissues dominated by 5p-guides more susceptible to DICER1 mutations? The cells used in this study are 3p-dominated; ∼40% of the total miRNA content of PA-1 cells derived from the miR-302a-3p and miR-302b-3p miRNAs. Given that 3p guides are not affected, cells with 3p-dominant profile could experience less severe impacts of DICER1 inactivation. Conversely, cell types and tissues that are 5p-dominated will likely experience a more dramatic change in their miRNA repertoire upon DICER1 inactivation.

In addition to miRNA function, we have explored the relevance of less well-characterised functions of DICER1 in the context of DICER1-synrome mutations. DICER1 is known to play roles in maintaining genome stability and ensuring proper chromosome segregation^5,6,9–11^, and misfunction in both can lead to aneuploidy. Our results showed, however, that endonucleolytic inactivation of DICER1 did not result in the types of chromosomal abnormalities reported in cells lacking DICER1. Unfortunately, we were not able to obtain full knockouts of *DICER1* in PA-1 cells to confirm these findings in our cellular system. It is possible that the complete inactivation of *DICER1* is lethal in PA-1 cells, as previously observed in human embryonic stem cells^26^. Nevertheless, we did obtain *DICER1^+/-^* cells, and interestingly, these cells did not show altered miRNA expression, suggesting that human *DICER1* is not haploinsufficient, as previously demonstrated for mouse *Dicer*^47^. The observation that human *DICER1* is not haploinsufficient may explain why germline carriers, where only one allele is inactivated, are viable, and they do not develop the disease until the second mutation occurs.

Previously, mouse *Dicer* had been implicated in the repression of transposable elements (TEs)^11,16^. We confirmed these findings in the context of human DICER1 endonucleolytic inactivation, showing that DICER1 represses both RNA retrotransposons and DNA transposons. We hypothesise that some of this repression may be driven by the production of small RNAs or endo-siRNAs. Typically, endo-siRNAs are generated by DICER1 cleavage of endogenous RNA substrates, including double-stranded RNA (dsRNA). This activity is typically observed only in pluripotent embryonic stem cells and oocytes ^19,20,48^. Interestingly, these two cell types, as well as embryonic teratocarcinoma cells, are devoid of the type I interferon (IFN) response, meaning they cannot produce IFN in response to cytoplasmic dsRNA accumulation, in contrast to somatic cell types (reviewed in^4^). Notably, PA-1 are derived from an ovarian embryonic teratocarcinoma, suggesting their IFN response is attenuated and that they can tolerate a certain level of endogenous dsRNA. In line with a potential role for endo-siRNAs in the control of TEs, we observed that DICER1 mutations affect the association of antisense small RNAs against the most upregulated LTR7 subfamily with AGO2. This regulation seems to be miRNA-independent, since cells lacking all canonical miRNAs do not display upregulation of LTR7 – HERVH elements, confirming previous findings for DGCR8-deficient human embryonic stem cells ^49^.

Although we attempted to assess the dysregulation of LTR7-HERVH expression in DICER1-mutated TCGA datasets, this was, unfortunately, not feasible due to the paucity of tumour-derived RNA-high throughput sequencing datasets with paired-end sequencing. However, HERVH reactivation has been observed in both liver metastases and primary colorectal cancers^50^. Although the role that HERVH plays in cancer development, the immune status, and its value as a predictive prognostic marker is still to be investigated.

The upregulation of LTR7 in DICER1 mutant cells has two potentially important consequences for our understanding of DICER1-syndrome. First, LTR7-HERVH, particularly the LTR7up1/up2 subfamilies, are the most highly expressed copies in human embryonic stem cells and are essential for the maintenance of pluripotency^51–54^. The parallels between pluripotent embryonic cell and cancer cells are striking: both undergo EMT transitions and require high self-renewal capacity^55^. Furthermore, the reactivation of LTR7 elements in cancer cells can lead to the generation of novel transcripts. For example, the activity of the LTR7 promoter in human embryonic stem cells results in the production of novel chimeric transcripts^53,56^. In the context of cancer, this property could be exploited to develop T-cell therapies against neo-antigens. For instance, *env* HERVH-encoded proteins have been used to train cytotoxic T cells to target colorectal carcinoma cells^57^. Additionally, the reactivation of LTRs can also result in the expression of oncogenes, in a process called onco-exaptation^58^. Exploring these dynamics in the context of DICER1 mutations could revolutionise our understanding of these tumours and their potential treatments.

## Methods

### Cell lines

Human PA-1 cells were cultured in Minimum Essential Medium (MEM) with GlutaMAX (Gibco) supplemented with 20% heat-inactivated fetal bovine serum (FBS,Gibco), 50ug/ml Gentamicin (Gibco) and 0.1mM MEM Non-Essential Amino Acids (Gibco) at 37°C with 5% CO_2_.

### Generation of CRISPR/CAS9 targeted clones

CRISPR RNA guides (crRNAs) were designed to target exon 21 or exon 24 close to the respective amino acid residues S1344 or D1709 in the RNase III a or b domains of DICER1 using the Cas9 design option from CRISPR Design Tool (https://chopchop.cbu.uib.no/) along with repair templates with the respective point mutations and silent mutations to remove PAM sites and introduction of unique restriction sites to aid screening to identify targeted clones.

crRNA-S1344 (5’-CACTTACCCTGATGCGCATG-3’), crRNA-D1709 (5’ AGTCCAAAATCGCATCTCCC-3’)

Repair template-S1344L

(5’CTGCAAACCACTTTCAGGCACACTGAATAATTAACTGCTCAAAATAAAAAAATCATCTCTTACCTTTTTGCTTCTCATATATAAAAGGCGCCCTTCATGCGCATCAGGGTAAGTGCAAAATAGATATGTGG-3’).

Repair template-D1709N

(5’CTTCGGATCCCCTCAGATTGTTACCAGCGCTTAGAATTTCTGGGAAATGCAATATTGGACTACCTCATAACCAAGCACCTTTATGAAGACCCGCGGCAGCACTCCCCGGGGGTCCTGACAGACCTGCGGTCTGCCCTGG-3’)

Equimolar amount of either crRNA-S1344 or crRNA-D1709 (245pmol, IDT) where incubated with tracrRNA (245 pmol, IDT) in 25ul nuclease-free buffer at 95°C for 5 mins and annealed by allowing to cool to room temperature (RT). These guides were then incubated with 25 µg Alt-R® S.p. Cas9 Nuclease V3 (IDT) in 22.5 µl buffer T (Neon Transfection System, ThermoFisher Scientific) for 20 minutes at 37°C prior to electroporation. PA-1 cells (1.2×10^6^ per transfection) were resuspended on 80 µl of Buffer T (Neon Transfection Systems), the Cas9 */* gRNA mixture and the corresponding repair template (300 pmol, IDT) were added to each tube. Electroporation was performed with three pulses of 1600 V and 10 ms. Cells were transferred to T25cm flask overnight before isolating single cell clones by limiting dilution in 96-well plates. The Platinum Direct PCR Universal Mastermix was used to screen for edited clones as per the manufacturer’s instructions using the following PCR primer pairs.

S1344L-F1 5’GAAATACCCGTGCAACCAA 3’

S1344L-R1 5’ TGTAAGCCAAGACGTACCCTC 3’

D1709N-F1 5’ CGGCCCTGGTTTCACTTAA 3’

D1709N-R1 5’ GCAGACAAGTTCAAGAGGCC 3’

The resulting PCR product (uncut) or diagnostic restriction digest was performed prior to running on 1.5% agarose gel and visualisation to identify targeted clones. The PCR product from targeted clones where either directly sequenced using Nanopore sequencing (Plasmidsarus) or cloned into pGEM-T Easy vector (Promega) and confirmed by Sanger sequencing. The homozygous point mutant *DICER1*^S1344L/S1344L^, *DICER1*^D1709N/D1709N^ and the compound heterozygous mutant *DICER1*^D1709N/FS^ were generated with a single round of targeting. To generate the *DICER1^S^*^1344^*^L/FS^* compound heterozygous mutant a second round of targeted was carried out using a heterozygous cell line *DICER1^S^*^1344^*^L/+^* which harboured the S1344L point mutation on one allele and PAM site silent mutation and the other allele was wild type.

### Metaphase Spreads

For karyotype analysis, each cell line was grown to approximately 80% confluency in a T75cm flask and treated for 30 min at 37°C with colcemid (KaryoMAX Colcemid, Gibco), trypsinised and cells ruptured using a hypotonic solution (0.075M KCl). The cell nuclei were then fixed using a 3:1 methanol/acetic acid solution and dropped on microscope slides before counting metaphase chromosomes (20 metaphase spreads per cell line were counted).

### Western blotting

Total cell lysates were prepared by lysing cell pellets in RIPA buffer supplemented with cOmplete EDTA-free protease inhibitor cocktail tablets (Roche), as per manufacturer’s instructions and centrifugation (13,000g for 20 minutes at 4°C) to clarify the extracts. The BCA assay (Pierce) was used to determine protein concentration and samples stored at-80°C until required. Proteins were separated on NuPage Novex Bis-Tris 4-20% Tris-Glycine protein gels (Invitrogen) and transferred to Immobilin-FL PVDF membrane (Sigma) by wet transfer. Membranes were blocked for 1hr at RT with 5% milk powder in TBS-T (0.1% Tween-20, Sigma) and incubated with primary antibodies overnight at 4°C. Primary antibodies used were mouse anti-DICER1 (1:1000, Clone 13D6, Cat # AB13D6, Abcam), rabbit anti-AGO2 (1:1000, Clone C34C6, Cat # 2897S, Cell Signalling) and mouse anti-Tubulin (1:4000 CP06, Millipore). Membranes were washed in TBS-T and incubated with secondary antibodies (Goat anti-rabbit IgG [H+L] or Goat anti-mouse IgG [H+L] for 1hr at RT and washed again before visualisation of protein bands using Odyssey CLx Scanner (LiCor). Protein bands were quantified using ImageJ (v1.53q) software and expression levels calculated normalized to tubulin.

### Growth rate /Proliferation assays

Cell growth rates were measured by seeding 1×10^5^ cells in triplicate in 6 well dish (day 0) and cell counts taken on the indicated days (3, 6, 9, 12 days) using a haemocytometer and cell counter before re-seeding.

### Metabolic activity assay

Metabolic activity was measured over time in the different cell lines by seeding cells in a 96 well plate (2×10^3^ cells per well) and measuring ATP levels using the CellTiter-Glo assay (Promega) at 7 time points (0,6,24,30,48,54,72hrs). Luminescence was recorded using Varioskan Flash (ThermoFisher Scientific) with an integration time of 1000ms per well.

### Colony formation assay

Triplicate 6-well dishes were seeded (5×10^2^ cells/well) and grown for 7 days, then washed with PBS, fixed with 70% ethanol and stained with 0.1% Crystal Violet (Sigma) before washing in water to remove background staining. Celigo image cytometer (Nexcelom Bioscience) was used to scan crystal violet-stained colonies in the brightfield channel. Software version 5.6.0.0 was used to calculate the number and size of colonies using the “Colony” application.

### Wound healing assay

Cells were seeded in triplicate in 6 well dishes to give a confluent monolayer the following day. A scratch was made in the centre of each well using a 20 μl plastic tip, the wound was gently washed with PBS to remove residual detached cells and fresh growth media added. Images were take using EVOS M7000 (x2 Objective AMEP431, transwell) at 0 h, 12 h, 24 h and 36 h post scratch. Quantification of the wound closure was measured using Image J software.

### Immunoprecipitation (IP)

Protein G beads (Invitrogen) were prepared by conjugating 50ul beads to 5 ug of anti-rat AGO2 antibody (Clone 11A9, Cat # MABE 253 Millipore) or rat IgG control per sample. The beads were equilibrated in lysis buffer (50 mM Tris-HCl pH 7.8, 300 mM NaCl, 1% Triton X-100, 5 mM EDTA, 10% Glycerol) prior to the addition of total cell lysates and incubated at 4°C overnight with rotation. The beads were washed as follows: Low Salt (LS)-wash (50 mM Tris-HCl pH 7.8, 300 mM NaCl, 5 mM MgCl_2_, 0.5% Triton X-100, 2.5% glycerol), twice and High Salt (HS)-wash (50 mM Tris-HCl pH 7.8, 800 mM NaCl, 5 mM MgCl_2_, 0.5% Triton X-100, 2.5% glycerol). For downstream protein analysis, the IP-protein fraction was eluted in LDS sample buffer (Life Technologies) containing 100 mM DTT (Sigma) at 70°C for 10 mins with shaking. For downstream RNA analysis, the IP-RNA fraction was resuspended in QIazol (700 µl) and RNA extracted using the miRNeasy mini kit (QIAGEN) following manufacturer’s instructions.

### Small RNA high throughput sequencing and analysis

RNA was extracted from the each of the five cell lines in triplicate using the miRNeasy mini kit for both input (total cell extracts) or AGO2-IPs. RNA recovered was quantified using the Qubit 2.0 Fluorometer (ThermoFisher Scientific, Q32866) and the quality assessed on the Agilent 2100 Electrophoresis Bioanalyser Instrument (Agilent, G2939AA). Library preparation and sequencing was carried out by the Clinical Research Facility, Edinburgh. For small RNA sequencing libraries, 500 ng of total RNA was used from each of the samples and libraries prepared using the QIAseq miRNA library kit 96 (QIAGEN, Cat #331505) and the QIAseq miRNA 96 Index Kit IL UDI-E (Cat #331655) according to the manufacturer’s protocols. Purified libraries were then amplified for 13 cycles of PCR with unique 10bp dual index (UDI) sequences to allow multiplexing on a single sequencing run. Amplified, indexed libraries were purified and size selected with QMN beads to enrich for library fragments containing miRNAs. Sequencing was performed on the NextSeq 2000 platform (Illumina Inc, #20038897) using NextSeq 1000/2000 P1 Reagents (100 Cycles) (#20074933). Sequencing was single end 1×75bp to allow coverage of the UMI sequence.

Small RNA-Seq data were processed as follows. FastQC and MultiQC were used to evaluate the quality of the raw reads (https://www.bioinformatics.babraham.ac.uk/projects/fastqc/)^59^.

Reaper was used to remove the 3’ adapter sequences, and distinct sequences were collapsed using tally, selecting those between 20-25nt, and keeping track of the counts of each read^60^. Unique reads were mapped against the human genome (GRCh38), using bowtie1 v1.3.1^61^ allowing 1 mismatch and reporting all possible locations (-v 1 --all --best –strata). A python script was then used to distribute each read’s counts evenly between all mapping locations. The remaining analyses were conducted within the R environment for statistical computing (https://www.r-project.org/). FeatureCounts was used to quantify overlapping reads^62^. Differential expression tests were performed within the edgeR framework, with quasi-dispersion estimates, and using glmTreat to test against a minimum fold-change threshold of 2 and a Benjamini-Hochberg corrected FDR < 0.01^63^. Normalisation factors were calculated manually using tRNAs as the reference annotation. Plotting used a combination of base R, ggplot2, ggside and ggrepel (https://ggplot2-book.org/). A custom genome annotation was developed, compiling protein-coding, lncRNA and tRNA annotation from Gencode r49^64^, microRNAs from miRBase r22.1^65^ and MirGeneDB r3.0^66^, mirtrons from mirtronDB v2^67^, and repeats as described under “Transposable elements analysis” below. Care was taken to remove redundancy due to overlaps, giving preference to known microRNA annotation.

### Modelling of DICER1 mutations

Coordinates from human DICER1 bound to pre-let7 (PDBid 7WX2^31^) were mutated *in silico*, using the mutation commands in Coot 0.9.6^68^, selecting a common rotomer for each mutant. For D1709L the rotomer most closely matching the sidechain in the experimental structure was used. Images of the mutant residues superimposed on the native structure were generated using Pymol 3.1.6.1 (https://www.pymol.org).

### Total RNA high throughput sequencing and analysis

Libraries for total RNA sequencing were prepared using the NEBNext Ultra 2 Directional RNA library prep kit (Illumina, E7760) and the NEBNext rRNA Depletion kit (Human/Mouse/Rat) (Illumina, E6310) according to the provided protocol. Sequencing was performed on the NextSeq 2000 platform (Illumina, 20038897) using NextSeq 2000 P3 Reagents (200 Cycles, 2×100bp) (20040559).

Total RNA-sequencings runs were processed with the following pipeline. Gene based read counts and TPMs were retrieved for each sample after mapping over human genome GRCh38/hg38 using Gencode39 annotation (Ensembl based) with STAR v2.7.9a^69^ with default parameters. Statistical inference of regulated genes of one comparison of grouped DICER1 mutants vs WT samples were calculated with the DESeq2 v1.34^70^ R library by setting the contrast parameter and weighted equally to the log2 fold changes of the four DICER1 mutants. Differentially regulated genes were retrieved with an FDR cut-off of 0.05 obtained with the Benjamini & Hochberg correction.

### Total cell proteomics

Total protein cell extracts were prepared by lysing cells in ice-cold RIPA buffer (50 mM Tris/HCl pH 7.4, 150 mM NaCl, 1% Triton-X100, 0.5% Sodium Deoxycholate, 0.1% SDS supplemented with cOmplete protease inhibitor cocktail tablets - Roche). Lysates were incubated on ice for 20 minutes. Insoluble cell debris was pelleted by centrifuged at 13,000g for 20 minutes at 4°C and the clarified total cell protein supernatant was saved as total cell extracts. Protein concentration was measured using BCA protein assay (ThermoFisher Scientific, 23225) as per manufacturer’s instructions. Four replicates were generated for each of the cell lines. For each individual replicate, proteins were purified using the SP3 magnetic bead method with a 1:1 mixture of Sera-Mag E3 and E7 beads. Protein samples were reduced with 10 mM dithiothreitol (DTT) for 30 min at 56 °C and alkylated with 50 mM iodoacetamide (IAA) for 30 min in the dark at room temperature, followed by quenching with 10 mM DTT. Beads were added at a ratio of 10 µg beads per 1 µg protein, and protein binding was induced by adding ethanol to a final concentration of 50%. Samples were mixed and incubated briefly for capture, followed by three washes with 80% ethanol and brief air-drying to remove residual solvent. Bound proteins were digested overnight at 37 °C in 50 mM ammonium bicarbonate using trypsin at a 1:50 enzyme-to-protein ratio (w/w). Digestion was stopped by acidification with trifluoroacetic acid (TFA), and peptides were captured on pre-equilibrated C18 StageTips. StageTips were washed to remove contaminants and stored at −20 °C, as described in^71^. Peptides were eluted in 40 μL of 80% acetonitrile (Fisher Scientific) in 0.1% TFA and concentrated down to 1 μL by vacuum centrifugation (Concentrator 5301, Eppendorf, UK). The peptide sample was then prepared for LC-MS/MS analysis by diluting it to 5 μL by 0.1% TFA.

LC-MS analyses were performed on an Orbitrap Exploris 480™ on Data Independent Acquisition (DIA), coupled on-line, to an Ultimate 3000 HPLC (Dionex, Thermo Fisher Scientific, UK). Peptides were separated on a 50 cm (2 µm particle size) EASY-Spray column (Thermo Scientific, UK), which was assembled on an EASY-Spray source (Thermo Scientific, UK) and operated constantly at 50°C. Mobile phase A consisted of 0.1% formic acid in LC-MS grade water and mobile phase B consisted of 80% acetonitrile and 0.1% formic acid. Peptides were loaded onto the column at a flow rate of 0.3 μL min^-1^ and eluted at a flow rate of 0.25 μL min^-1^ according to the following gradient: 2 to 40% mobile phase B in 150 min and then to 95% in 11 min. Mobile phase B was retained at 95% for 5 min and returned back to 2% a minute after until the end of the run (190 min).

MS1 scans were recorded at 120,000 resolution (scan range 350-1650 m/z) with an ion target of 5.0e6, and injection time of 20ms. MS2 DIA was performed in the orbitrap at 30,000 resolution with a scan range of 350-1200 m/z, maximum injection time of 55ms and AGC target of 3.0E6 ions. We used HCD fragmentation^72^ with stepped collision energies of 25.5, 27 and 30. We used variable isolation windows throughout the scan range ranging from 10.5 to 50.5 m/z. Shorter isolation windows (10.5-18.5 m/z) were applied from 400-800 m/z and then gradually increased to 50.5 m/z until the end of the scan range. The default charge state was set to 3. Data for both survey and MS/MS scans were acquired in profile mode.

The DIA-NN software platform^73^ version 1.9.2. was used to process the DIA raw files and search was conducted against the canonical *Homo sapiens* database (Uniprot, released August 2022). Precursor ion generation was based on the chosen protein database (automatically generated spectral library) with deep-learning based spectra, retention time and IMs prediction. Digestion mode was set to specific with trypsin allowing maximum of two missed cleavages. Carbamidomethylation of cysteine was set as fixed modification. Oxidation of methionine, and acetylation of the N-terminus were set as variable modifications. The parameters for peptide length range, precursor charge range, precursor m/z range and fragment ion m/z range as well as other software parameters were used with their default values. The precursor FDR was set to 1%.

Differential protein abundance was performed using Perseus (proteinGroups) outputs. Keratins were filtered out as contaminants, proteins missing values (N/A) across all compared samples were also excluded. For relative quantification, protein row intensities were normalised and transformed using the R-package Variance Stabilising Normalisation, VSN (3.74.0)^74^. Missing value Imputation was performed by determining the global minimum value, and imputation was performed only for proteins undetected in all replicates in one experimental condition, while present in the other condition (at least in 2 replicates, ON/OFF). The R package limma (3.62.2)^75^ was used for statistical analysis of the processed protein intensities, applying the empirical Bayesian method moderated t-test with p-values adjusted for multiple testing using the Benjamini-Hochberg method.

### Transposable elements analysis

Transposable element annotation for hg38 was obtained from the UCSC Genome Browser. A defragmented annotation was obtained by merging individual TE fragments into whole copies using Onecodetofindthemall.pl^76^, and defragmented copies were renamed according to their position and structure. This defragmented annotation was used to perform locus-specific quantification. To specifically analyse the expression of LTR7 subfamilies, we used the recent annotation by Carter et al ^35^.

TE expression both at family and individual locus level was quantified using TEcount and TElocal tools from TEtranscript toolkit, respectively^77^. First, defragmented TE annotation was indexed for TElocal using script TElocal_indexer provided by TEtranscript developers, and sequencing reads were aligned with STAR v2.7.11b^69^ using parameters recommended by authors in^77^. Both TEcount and TElocal distributed multimapped reads among all alignments.

Differential expression analysis was carried out using DESeq2^70^, estimating size factors considering only genes and applying lfcShrink function to shrink log2 fold changes. Volcano and violin plots were generated using in-house R scripts with ggplot2.

For annotation of LTR7-derived small RNAs, size selection was performed with pullseq (1.0.2; https://github.com/bcthomas/pullseq), extracting sequences of 20-25 nucleotides in length.

Total (non-unique) reads were aligned to the LTR7 consensus sequences with bowtie1, as above. Samtools (1.22.1)^78^ was used to sort and index mapped sequences. Aligned sequences were normalised as counts per 100 million sequence reads, and split into plus or minus strands via BEDtools (2.27.1)^79^. Sequence pileups were visualised using IGV^80^.

### TCGA-UCEC microRNA analysis

Quantification for isoform microRNA expression was retrieved for TCGA-UCEC tumour samples using TCGAbiolinks package^81^ and downloaded with the GDC Data Transfer Tool. microRNA samples were classified as DICER1 mutated hotspots (as described in ^24^) and non-DICER1 samples. Non-DICER1 samples were further filtered to remove samples with mutations in genes involved in microRNA biogenesis (DICER1, DROSHA, DGCR8 and AGO family genes).

MicroRNAs were assigned to 5p or 3p group according to their mirBase v21 name prefix. Fragments annotated as precursor, stemloop, or unannotated were excluded. Differential expression analysis of DICER1 tumour samples (n=14) versus non-DICER1 samples (n=367) was carried out with DESeq2. microRNAs were considered dysregulated if p-adj < 0.05.

### miRNA predicted targets and comparison to transcriptomic/proteomic data

Predictions for all human miRNA targets were obtained from Targetscan r8.0^82^, selecting the cumulative weighted context++ scores (CWCS) from the “Summary Counts for all predictions” file. Target genes were mapped to the transcriptomic and proteomic experiments using the official Gene Symbol. To understand if predicted targets were changing as expected, we focused on the 10 most up/down-regulated miRNAs from the D1709N or S1344L mutants vs WT sRNA-Seq data (fold-change > 10, FDR < 0.05 and sorted by average CPM). For each of these groups of miRNAs we chose the top 500 targets (according to CWCS) that were also measured in the transcriptomic or proteomic experiment. Assuming that the effect of a gene targeted by up-regulated as well as down-regulated miRNAs would be difficult to predict, we excluded genes that were also within the top 1000 predicted targets of miRNAs changing in the opposite direction. The remaining genes (<500) were analysed by plotting their logFC cumulative distribution and by performing a one-tailed Wilcoxon Rank Sum test against the genes that were not in any of the predicted target groups. The alternative hypothesis for up(down)-regulated miRNAs was that targets will be down(up)-regulated.

## Acknowledgements

This work was funded by the Wellcome Trust grants (221737/Z/20/Z) and (107665/Z/15/Z) and the Leverhulme Trust (RPG-2020-355) to S.M. F.M. and K.K. were funded by Darwin Trust fellowships. The S.R.H. laboratory work was supported by grant CNS2023-145402 funded by MICIU/AEI/10.13039/501100011033 and by “European Union NextGeneration EU/PRTR”. This work was also supported by funding for the Wellcome Discovery Research Platform for Hidden Cell Biology (0226791), and we gratefully acknowledge support from the Proteomics core. We also thank Dr. Petra Schneider (Institute of Ecology and Evolution, University of Edinburgh) for help with the statistical analysis, David Wright (Institute of Immunology and Infection Research-IIIR, University of Edinburgh-UoE) for help with CRISPR/Cas9 design, and Dr. Jessica L. Walters (IIIR, UoE) for help with analysis of the CRISPR-edited clones. We thank the computer resources (Picasso Supercomputer), technical expertise and assistance provided by the SCBI (Supercomputing and Bioinformatics) Center of the University of Malaga.

## Contributions

S.M. and K.G. conceived the project and wrote the manuscript with the help from all authors. K.G. performed most experiments with PA-1 cells, with the help from J.W. and J.W. N.B. performed the bioinformatic analyses of the total RNA-sequencing dataset, and initial analysis of miRNA expression. F.M and K.G. performed the proteomic analysis with support from the Proteomics Core facility, at the Wellcome Discovery Research Platform for Hidden Cell Biology. P.G.M and G.P. performed the TE analysis under the supervision of S.R.H. S.B. performed chromosome karyotyping. A.C. performed the *in silico* mutagenesis of DICER1. C.A.G and K.K performed the small RNA and AGO2 IP bioinformatic analysis.

## References

1. Medley, J. C., Panzade, G. & Zinovyeva, A. Y. microRNA strand selection: Unwinding the rules. Wiley Interdiscip. Rev. RNA 12, e1627 (2020).

2. Suzuki, H. I. et al. Small-RNA asymmetry is directly driven by mammalian Argonautes. Nat. Struct. Mol. Biol. 22, 512–521 (2015).

3. Chen, L. et al. miRNA arm switching identifies novel tumour biomarkers. eBioMedicine 38, 37–46 (2018).

4. Mueller, F., Witteveldt, J. & Macias, S. Antiviral Defence Mechanisms during Early Mammalian Development. Viruses 16, 173 (2024).

5. Pek, J. W. & Kai, T. DEAD-box RNA helicase Belle/DDX3 and the RNA interference pathway promote mitotic chromosome segregation. Proc. Natl. Acad. Sci. 108, 12007–12012 (2011).

6. Harfe, B. D., McManus, M. T., Mansfield, J. H., Hornstein, E. & Tabin, C. J. The RNaseIII enzyme Dicer is required for morphogenesis but not patterning of the vertebrate limb. Proc. Natl. Acad. Sci. 102, 10898–10903 (2005).

7. Fukagawa, T. et al. Dicer is essential for formation of the heterochromatin structure in vertebrate cells. Nat. Cell Biol. 6, 784–791 (2004).

8. Murchison, E. P., Partridge, J. F., Tam, O. H., Cheloufi, S. & Hannon, G. J. Characterization of Dicer-deficient murine embryonic stem cells. Proc. Natl. Acad. Sci. 102, 12135–12140 (2005).

9. Huang, C., Wang, X., Liu, X., Cao, S. & Shan, G. RNAi pathway participates in chromosome segregation in mammalian cells. Cell Discov. 1, 15029 (2015).

10. Yadav, R. P., Mäkelä, J.-A., Hyssälä, H., Cisneros-Montalvo, S. & Kotaja, N. DICER regulates the expression of major satellite repeat transcripts and meiotic chromosome segregation during spermatogenesis. Nucleic Acids Res. 48, 7135–7153 (2020).

11. Gutbrod, M. J. et al. Dicer promotes genome stability via the bromodomain transcriptional co-activator BRD4. Nat. Commun. 13, 1001 (2022).

12. Cornec, A. & Poirier, E. Z. Interplay between RNA interference and transposable elements in mammals. Front. Immunol. 14, (2023).

13. Heras, S. R., Macias, S., Cáceres, J. F. & Garcia-Perez, J. L. Control of mammalian retrotransposons by cellular RNA processing activities. Mob. Genet. Elem. 4, e28439 (2014).

14. Svoboda, P. et al. RNAi and expression of retrotransposons MuERV-L and IAP in preimplantation mouse embryos. Dev. Biol. 269, 276–285 (2004).

15. Kanellopoulou, C. et al. Dicer-deficient mouse embryonic stem cells are defective in differentiation and centromeric silencing. Genes Dev. 19, 489–501 (2005).

16. Bodak, M., Cirera-Salinas, D., Yu, J., Ngondo, R. P. & Ciaudo, C. Dicer, a new regulator of pluripotency exit and LINE-1 elements in mouse embryonic stem cells. FEBS Open Bio 7, 204–220 (2017).

17. Berrens, R. V. et al. An endosiRNA-Based Repression Mechanism Counteracts Transposon Activation during Global DNA Demethylation in Embryonic Stem Cells. Cell Stem Cell 21, 694–703.e7 (2017).

18. Yang, N. & Kazazian, H. H. L1 retrotransposition is suppressed by endogenously encoded small interfering RNAs in human cultured cells. Nat. Struct. Mol. Biol. 13, 763–771 (2006).

19. Watanabe, T. et al. Identification and characterization of two novel classes of small RNAs in the mouse germline: retrotransposon-derived siRNAs in oocytes and germline small RNAs in testes. Genes Dev. 20, 1732–1743 (2006).

20. Babiarz, J. E., Ruby, J. G., Wang, Y., Bartel, D. P. & Blelloch, R. Mouse ES cells express endogenous shRNAs, siRNAs, and other Microprocessor-independent, Dicer-dependent small RNAs. Genes Dev. 22, 2773–2785 (2008).

21. Calabrese, J. M., Seila, A. C., Yeo, G. W. & Sharp, P. A. RNA sequence analysis defines Dicer’s role in mouse embryonic stem cells. Proc. Natl. Acad. Sci. 104, 18097–18102 (2007).

22. Foulkes, W. D., Priest, J. R. & Duchaine, T. F. DICER1: mutations, microRNAs and mechanisms. Nat. Rev. Cancer 14, 662–672 (2014).

23. Kim, H., Lee, Y.-Y. & Kim, V. N. The biogenesis and regulation of animal microRNAs. Nat. Rev. Mol. Cell Biol. 26, 276–296 (2025).

24. Vedanayagam, J. et al. Cancer-associated mutations in DICER1 RNase IIIa and IIIb domains exert similar effects on miRNA biogenesis. Nat. Commun. 10, 3682 (2019).

25. Bernstein, E. et al. Dicer is essential for mouse development. Nat. Genet. 35, 215–217 (2003).

26. Teijeiro, V. et al. DICER1 Is Essential for Self-Renewal of Human Embryonic Stem Cells. Stem Cell Rep. 11, 616–625 (2018).

27. Pastushenko, I. & Blanpain, C. EMT Transition States during Tumor Progression and Metastasis. Trends Cell Biol. 29, 212–226 (2019).

28. Youssef, K. K. et al. Two distinct epithelial-to-mesenchymal transition programs control invasion and inflammation in segregated tumor cell populations. *Nat*. Cancer 5, 1660–1680 (2024).

29. Choucair, K. et al. Targeting KRAS mutations: orchestrating cancer evolution and therapeutic challenges. Signal Transduct. Target. Ther. 10, 385 (2025).

30. Liberti, M. V. & Locasale, J. W. The Warburg Effect: How Does it Benefit Cancer Cells? Trends Biochem. Sci. 41, 211–218 (2016).

31. Lee, Y.-Y., Lee, H., Kim, H., Kim, V. N. & Roh, S.-H. Structure of the human DICER–pre-miRNA complex in a dicing state. Nature 615, 331–338 (2023).

32. Malagobadan, S. et al. DICER1 hotspot mutation induces 3p microRNA gain of function via Argonaute strand switch. Nat. Struct. Mol. Biol. 1–11 (2025) doi:10.1038/s41594-025-01671-w.

33. Jee, D. et al. Human DICER1 hotspot mutation induces both loss and gain of miRNA function. Nat. Struct. Mol. Biol. 1–11 (2025) doi:10.1038/s41594-025-01701-7.

34. Gázquez-Gutiérrez, A. et al. Control of retrotransposon-driven activation of the interferon response by the double-stranded RNA binding protein DGCR8. 2025.05.28.656609 Preprint at 10.1101/2025.05.28.656609 (2025).

35. Carter, T. A. et al. Mosaic cis-regulatory evolution drives transcriptional partitioning of HERVH endogenous retrovirus in the human embryo. eLife 11, e76257 (2022).

36. Römer, C., Singh, M., Hurst, L. D. & Izsvák, Z. How to tame an endogenous retrovirus: HERVH and the evolution of human pluripotency. Curr. Opin. Virol. 25, 49–58 (2017).

37. Wu, M. K. et al. Biallelic DICER1 mutations occur in Wilms tumours. J. Pathol. 230, 154–164 (2013).

38. Wegert, J. et al. Mutations in the SIX1/2 Pathway and the DROSHA/DGCR8 miRNA Microprocessor Complex Underlie High-Risk Blastemal Type Wilms Tumors. Cancer Cell 27, 298–311 (2015).

39. Torrezan, G. T. et al. Recurrent somatic mutation in DROSHA induces microRNA profile changes in Wilms tumour. Nat. Commun. 5, 4039 (2014).

40. Walz, A. L. et al. Recurrent DGCR8, DROSHA, and SIX Homeodomain Mutations in Favorable Histology Wilms Tumors. Cancer Cell 27, 286–297 (2015).

41. Tiburcio, P. D. B. et al. DROSHA Regulates Mesenchymal Gene Expression in Wilms Tumor. Mol. Cancer Res. 22, 711–720 (2024).

42. Rakheja, D. et al. Somatic mutations in DROSHA and DICER1 impair microRNA biogenesis through distinct mechanisms in Wilms tumours. Nat. Commun. 5, 4802 (2014).

43. Anglesio, M. et al. Cancer-associated somatic DICER1 hotspot mutations cause defective miRNA processing and reverse-strand expression bias to predominantly mature 3p strands through loss of 5p strand cleavage. J. Pathol. 229, 400–409 (2013).

44. Nadaf, J. et al. Molecular characterization of DICER1-mutated pituitary blastoma. Acta Neuropathol. (Berl.) 141, 929–944 (2021).

45. Ricarte-Filho, J. C. et al. DICER1 RNase IIIb domain mutations trigger widespread miRNA dysregulation and MAPK activation in pediatric thyroid cancer. Front. Endocrinol. 14, (2023).

46. Kommoss, F. K. F. et al. Genomic characterization of DICER1-associated neoplasms uncovers molecular classes. Nat. Commun. 14, 1677 (2023).

47. Murchison, E. P., Partridge, J. F., Tam, O. H., Cheloufi, S. & Hannon, G. J. Characterization of Dicer-deficient murine embryonic stem cells. Proc. Natl. Acad. Sci. 102, 12135–12140 (2005).

48. Flemr, M. et al. A Retrotransposon-Driven Dicer Isoform Directs Endogenous Small Interfering RNA Production in Mouse Oocytes. Cell 155, 807–816 (2013).

49. Colomer-Boronat, A. et al. DGCR8 haploinsufficiency leads to primate-specific RNA dysregulation and pluripotency defects. Nucleic Acids Res. 53, gkaf197 (2025).

50. Viot, J. et al. Deciphering human endogenous retrovirus expression in colorectal cancers: exploratory analysis regarding prognostic value in liver metastases. eBioMedicine 116, 105727 (2025).

51. Ohnuki, M. et al. Dynamic regulation of human endogenous retroviruses mediates factor-induced reprogramming and differentiation potential. Proc. Natl. Acad. Sci. 111, 12426–12431 (2014).

52. Lu, X. et al. The retrovirus HERVH is a long noncoding RNA required for human embryonic stem cell identity. Nat. Struct. Mol. Biol. 21, 423–425 (2014).

53. Wang, J. et al. Primate-specific endogenous retrovirus-driven transcription defines naive-like stem cells. Nature 516, 405–409 (2014).

54. Izsvák, Z., Ma, J., Singh, M. & Hurst, L. D. Co-option of an endogenous retrovirus (LTR7-HERVH) in early human embryogenesis: becoming useful and going unnoticed. Mob. DNA 16, 27 (2025).

55. Manzo, G. Similarities Between Embryo Development and Cancer Process Suggest New Strategies for Research and Therapy of Tumors: A New Point of View. Front. Cell Dev. Biol. 7, 20 (2019).

56. Wang, Y. et al. Endogenous miRNA Sponge lincRNA-RoR Regulates Oct4, Nanog, and Sox2 in Human Embryonic Stem Cell Self-Renewal. Dev. Cell 25, 69–80 (2013).

57. Mullins, C. S. & Linnebacher, M. Endogenous retrovirus sequences as a novel class of tumor-specific antigens: an example of HERV-H env encoding strong CTL epitopes. Cancer Immunol. Immunother. 61, 1093–1100 (2012).

58. Babaian, A. et al. Onco-exaptation of an endogenous retroviral LTR drives IRF5 expression in Hodgkin lymphoma. Oncogene 35, 2542–2546 (2016).

59. Ewels, P., Magnusson, M., Lundin, S. & Käller, M. MultiQC: summarize analysis results for multiple tools and samples in a single report. Bioinformatics 32, 3047–3048 (2016).

60. Davis, M. P. A., van Dongen, S., Abreu-Goodger, C., Bartonicek, N. & Enright, A. J. Kraken: A set of tools for quality control and analysis of high-throughput sequence data. Methods 63, 41–49 (2013).

61. Langmead, B., Trapnell, C., Pop, M. & Salzberg, S. L. Ultrafast and memory-efficient alignment of short DNA sequences to the human genome. Genome Biol. 10, R25 (2009).

62. Liao, Y., Smyth, G. K. & Shi, W. The R package Rsubread is easier, faster, cheaper and better for alignment and quantification of RNA sequencing reads. Nucleic Acids Res. 47, e47 (2019).

63. Chen, Y., Chen, L., Lun, A. T. L., Baldoni, P. L. & Smyth, G. K. edgeR v4: powerful differential analysis of sequencing data with expanded functionality and improved support for small counts and larger datasets. Nucleic Acids Res. 53, gkaf018 (2025).

64. Mudge, J. M. et al. GENCODE 2025: reference gene annotation for human and mouse. Nucleic Acids Res. 53, D966–D975 (2025).

65. Kozomara, A., Birgaoanu, M. & Griffiths-Jones, S. miRBase: from microRNA sequences to function. Nucleic Acids Res. 47, D155–D162 (2019).

66. Clarke, A. W. et al. MirGeneDB 3.0: improved taxonomic sampling, uniform nomenclature of novel conserved microRNA families and updated covariance models. Nucleic Acids Res. 53, D116–D128 (2025).

67. Da Fonseca, B. H. R., Domingues, D. S. & Paschoal, A. R. mirtronDB: a mirtron knowledge base. Bioinformatics 35, 3873–3874 (2019).

68. Emsley, P., Lohkamp, B., Scott, W. G. & Cowtan, K. Features and development of Coot. Acta Crystallogr. D Biol. Crystallogr. 66, 486–501 (2010).

69. Dobin, A. et al. STAR: ultrafast universal RNA-seq aligner. Bioinformatics 29, 15–21 (2013).

70. Love, M. I., Huber, W. & Anders, S. Moderated estimation of fold change and dispersion for RNA-seq data with DESeq2. Genome Biol. 15, 550 (2014).

71. Rappsilber, J., Mann, M. & Ishihama, Y. Protocol for micro-purification, enrichment, pre-fractionation and storage of peptides for proteomics using StageTips. Nat. Protoc. 2, 1896–1906 (2007).

72. Olsen, J. V. et al. Higher-energy C-trap dissociation for peptide modification analysis. Nat. Methods 4, 709–712 (2007).

73. Demichev, V., Messner, C. B., Vernardis, S. I., Lilley, K. S. & Ralser, M. DIA-NN: neural networks and interference correction enable deep proteome coverage in high throughput. Nat. Methods 17, 41–44 (2020).

74. Huber, W., von Heydebreck, A., Sültmann, H., Poustka, A. & Vingron, M. Variance stabilization applied to microarray data calibration and to the quantification of differential expression. Bioinformatics 18, S96–S104 (2002).

75. Ritchie, M. E. et al. limma powers differential expression analyses for RNA-sequencing and microarray studies. Nucleic Acids Res. 43, e47 (2015).

76. Bailly-Bechet, M., Haudry, A. & Lerat, E. “One code to find them all”: a perl tool to conveniently parse RepeatMasker output files. Mob. DNA 5, 13 (2014).

77. Jin, Y., Tam, O. H., Paniagua, E. & Hammell, M. TEtranscripts: a package for including transposable elements in differential expression analysis of RNA-seq datasets. Bioinforma. Oxf. Engl. 31, 3593–3599 (2015).

78. Danecek, P. et al. Twelve years of SAMtools and BCFtools. GigaScience 10, giab008 (2021).

79. Quinlan, A. R. & Hall, I. M. BEDTools: a flexible suite of utilities for comparing genomic features. Bioinformatics 26, 841–842 (2010).

80. Thorvaldsdóttir, H., Robinson, J. T. & Mesirov, J. P. Integrative Genomics Viewer (IGV): high-performance genomics data visualization and exploration. Brief. Bioinform. 14, 178–192 (2013).

81. Colaprico, A. et al. TCGAbiolinks: an R/Bioconductor package for integrative analysis of TCGA data. Nucleic Acids Res. 44, e71 (2016).

82. McGeary, S. E. et al. The biochemical basis of microRNA targeting efficacy. Science 366, eaav1741 (2019).

